# Human genetics and clinical aspects of neurodevelopmental disorders

**DOI:** 10.1101/000687

**Authors:** Gholson J. Lyon, Jason O’Rawe

## Introduction

> *“our incomplete studies do not permit actual classification; but it is better to leave things by themselves rather than to force them into classes which have their foundation only on paper” — Edouard Seguin* (Seguin, 1866)

> *“The fundamental mistake which vitiates all work based upon Mendel’s method is the neglect of ancestry, and the attempt to regard the whole effect upon offspring, produced by a particular parent, as due to the existence in the parent of particular structural characters; while the contradictory results obtained by those who have observed the offspring of parents apparently identical in certain characters show clearly enough that not only the parents themselves, but their race, that is their ancestry, must be taken into account before the result of pairing them can be predicted” — Walter Frank Raphael Weldon (Weldon,1902)*.

There are ~12 billion nucleotides in every cell of the human body, and there are ~25–100 trillion cells in each human body. Given somatic mosaicism, epigenetic changes and environmental differences, no two human beings are the same, particularly as there are only ~7 billion people on the planet. One of the next great challenges for studying human genetics will be to acknowledge and embrace complexity (Allchin, 2005; Bearn, 1993; Comfort, 2012; Grillo et al., 2013; Misteli, 2013; Radick, 2011; Sabin et al., 2013; Scriver, 2007; Tennessen etal., 2012; Terwilliger and Weiss, 2003; Weiss and Terwilliger, 2000). Every human *is* unique, and the study of human disease phenotypes (and phenotypes in general) will be greatly enriched by moving from a deterministic to a more stochastic/probabilistic model (Freund et al., 2013; Gigerenzer, 2002; Gigerenzer and Galesic, 2012; Gigerenzer et al., 2010; Kurz-Milcke et al., 2008; Sokal, 2012). The dichotomous distinction between ‘simple’ and ‘complex’ diseases is completely artificial, and we argue instead for a model that considers a spectrum of diseases that are variably manifesting in each person. The rapid adoption of whole genome sequencing (WGS) and the Internet-mediated networking of people promise to yield more insight into this century-old debate (Bateson and Mendel, 1902; Lyon and Segal, 2013; Lyon and Wang, 2012; Nielsen, 2012; Olby, 1989; Provine, 2001; Weldon, 1902). Comprehensive ancestry tracking and detailed family history data, when combined with WGS or at least cascade-carrier screening (McClaren et al., 2010), might eventually facilitate a degree of genetic prediction for some diseases in the context of their familial and ancestral etiologies. However, it is important to remain humble, as our current state of knowledge is not yet sufficient, and in principle, any number of nucleotides in the genome, if mutated or modified in a certain way and at a certain time and place, might influence some phenotype during embryogenesis or postnatal life (Batista and Chang, 2013; Cartault et al., 2012; Dickel et al., 2013; Hansen et al., 2013; Kapusta et al., 2013; Keller, 2010; Khoddami and Cairns, 2013; Ledford, 2013; Maxmen, 2013; Memczak et al., 2013; Mercer and Mattick, 2013; Miura et al., 2013; Moreau et al., 2013; Ning et al., 2013; Pennacchio et al., 2013; Perrat et al., 2013; Sabin et al., 2013; Salzman et al., 2012; Wilusz and Sharp, 2013).

In this chapter, we will traverse contemporary understandings of the genetic architecture of human disease, and explore the clinical implications of the current state of our knowledge. Many molecular models have been postulated as being important in genetic disease, and, despite our incomplete knowledge of the genetic workings of many diseases, significant progress has been made over the past 50 years. Many different classes of genetic mutations have been implicated as being involved in predisposition to certain diseases, and we are continually uncovering other means by which genetics plays an important role in human disease, such as with somatic genetic mosaicism. An explosion in the development of new biomedical techniques, molecular technologies, and analytical tools has enriched our knowledge of the many molecular bases of disease, underscored by the fact that we now exist in a world where each person can be characterized on the level of their ‘genome’, ‘transcriptome’ and ‘proteome’. We discuss these exciting new developments and the current applications of these technologies, their limitations, their implications for prenatal diagnosis and implantation genetics, as well as future prospects.

## Clinical classifications and the genetic architecture of disease

> *“Those who have given any attention to congenital mental lesions, must have been frequently puzzled how to arrange, in any satisfactory way, the different classes of this defect which may have come under their observation. Nor will the difficulty be lessened by an appeal to what has been written on the subject. The systems of classification are generally so vague and artificial, that, not only do they assist but feebly, in any mental arrangement of the phenomena represented, but they completely fail in exerting any practical influence on the subject.” —John Langdon Down* (Down, 1995)

As most clinicians know from experience, it is quite difficult to characterize the range of human experience in the two-dimensional world of the printed page, as we are attempting to do here. In addition, classifications can sometimes lead people to try to force round pegs into square holes, and so we are reluctant to further promulgate these classifications. Such classifications include terms such as: ‘Mendelian’, ‘complex disease’, ‘penetrance’, ‘expressivity’, ‘oligogenic’, and ‘polygenic’. For example, some have used the word ‘Mendelian’ to refer to a disease that appears to somehow be ‘caused’ by mutations in a single gene. As such, cystic fibrosis, Huntington’s disease, and Fragile X are all diseases that some people refer to as being ‘caused’ by mutations occurring in single genes. However, the expression of the phenotype within these diseases is extremely variable, depending in part on the exact mutations in each gene, and it is not at all clear that any mutation really and truly ‘causes’ any phenotype, at least not according to thoughtful definitions of causation that we are aware of (Fins, 2009; Hume and Selby-Bigge, 1896). For example, some children with certain mutations in *CFTR* may only have pancreatitis as a manifestation of cystic fibrosis, without any lung involvement (Corleto et al., 2010; Derikx and Drenth, 2010), and there is evidence that mutations in other genes in the genomes can have a modifying effect on the phenotype (Emond et al., 2012; Rosendahl et al., 2013). In the case of Huntington’s, there is extreme variability in the expression of the phenotype, both in time, period and scope of illness, and all of this is certainly modified substantially by the number of trinucleotide repeats (Orr and Zoghbi, 2007), genetic background (Tome et al., 2013) and environmental influences (Ciancarelli et al., 2013). Even in the case of whole chromosome disorders, such as Down Syndrome, there is ample evidence of substantial phenotypic expression differences, modified again by genetic background (Ackerman et al., 2012; Li et al., 2012), somatic mosaicism (Papavassiliou et al., 2009) and environmental influences (Dodd and Shields, 2005; Solomon, 2012), including synaptic and brain plasticity (Freund et al., 2013; Maffei, 2012; Maffei et al., 2012; Maffei and Turrigiano, 2008; Wang et al., 2013). The same is true for genomic deletion and duplication syndromes, such as velocardiofacial syndrome and other deletions (Guris et al., 2006; Iascone et al., 2002; Liao et al., 2004; McDonald-McGinn et al., 2013; Moreno-De-Luca et al., 2013a; Stalmans et al., 2003). And, of course, there is constant interaction of the environment with a person, both prenatally and postnatally. As just one example, cretinism is related to a lack of iodine in the mother’s diet, and there is incredibly variable expression of this illness based in part on the amount of iodine deficiency and how this interacts with fetal development (Zimmermann, 2012).

The words ‘penetrance’ and ‘expressivity’ can be defined as:

- *Penetrance*: The number of individuals in a population carrying a disease predisposing allele that are also categorically defined as being affected by the associated disease.
- *Expressivity*: The extremeness, or number of symptoms, in the presentation of a disease in the context of individuals who have the associated disease predisposing allele.”

Unfortunately, these two separate terms have led to a great deal of confusion in the field, and this sort of categorical thinking tends to miss complexity. Some use the word ‘penetrance’ when they really mean ‘expressivity’ of disease in any one person. As such, perhaps we should get rid of the two terms altogether and just discuss the expression of each trait in the context of a phenotypic spectrum, which is of course what led Walter Frank Raphael Weldon to establish the field of biometry (Jamieson and Radick, 2013; McIntyre, 2008; Sokal and Rohlf, 2012). Another way to express this point is to say that we have yet to characterize the full breadth of expression for virtually any mutation in humans, as we have not systematically sequenced or karyotyped any genetic alteration in thousands to millions of randomly selected people from a whole range of ethnic classes, i.e. clans (Bittles and Black, 2010; Lupski et al., 2011). There is an ongoing clash of worldviews, with some wanting to believe that single mutations predominately drive outcome while others are explicitly acknowledging the importance of substantial phenotypic modification via genetic background and/or environmental influence(s) (Beaudet, 2013; Bernal and Jirtle, 2010; Burga et al., 2011; Casanueva et al., 2012; Comfort, 2001, 2012; Dolinoy et al., 2006; Keller, 2010; Weinhouse et al., 2011). Some recent population-based sequencing efforts have shown the complexity of demonstrating how much any one genetic variant contributes to disease in any one particular individual, and we disagree with overly simplistic and artificial categorizations of mutations as “causative”, “pathogenic” or “non-pathogenic” (Andreasen et al., 2013a; Andreasen et al., 2013b; Refsgaard et al., 2012; Risgaard et al., 2013).

It is very likely that there will be a continuum of disease, given that the ‘effect size’ of any particular mutation will obviously vary according to genetic background and environment, as demonstrated repeatedly in model organisms (Bernal and Jirtle, 2010; Blount et al., 2012; Casanueva et al., 2012; Dolinoy et al., 2006; Greenspan, 2008, 2009, 2012; Holmes and Summers, 2006; Kendler and Greenspan, 2006; Meyer et al., 2012a; van Swinderen and Greenspan, 2005; Weinhouse et al., 2011). Thus, while a mutation associated with hemochromatosis or breast cancer might have high expression in one particular pedigree or clan, that same mutation may have very low expression in another pedigree, clan or group of unrelated people (Kohane et al., 2012). The reasons for variable expression can be myriad and are currently unknown in many instances; however, problems start to appear when scientists attempt to invoke a third allele as necessary and perhaps sufficient for the expression of any symptoms from within a typical disease. This disease model has been most clearly advocated for Bardet-Biedl Syndrome, in which the authors contend that some subjects have zero disease symptoms while possessing two autosomal recessive mutations in a known “disease gene”; the authors also show that some affected people have a mutation in another gene, i.e. a third allele, which they speculate is necessary and perhaps sufficient for expression of any symptoms of the disease (Eichers et al., 2004; Katsanis et al., 2001; Katsanis et al., 2002). However, this model has been challenged by others (Abu-Safieh et al., 2012; Laurier et al., 2006; Mykytyn et al., 2003; Nakane and Biesecker, 2005; Smaoui et al., 2006), and at least one group maintains that all people that they have studied with two autosomal recessive mutations have some manifestations of disease but with variable expression, i.e. one person might only have retinitis pigmentosa whereas another person might have the full-blown symptoms of Bardet-Biedl syndrome (Abu-Safieh et al., 2012). One wonders whether the debate about triallelism might really just be a semantic one due to problems with the phenotyping of ‘unaffected’ people, particularly if these people were not evaluated longitudinally. Detailed online longitudinal characterizations of all such reportedly ‘unaffected’ people could aid in documenting, with some degree of certainty, that these people did indeed have zero symptoms of Bardet-Biedl syndrome, as that would then be further proof that mutations are not deterministic in any way at all. Said another way, this would be demonstration of the enormous variability in expression for mutations that do contribute more to a phenotype in some people with their own genetic backgrounds and environmental differences, and this observation ought to have dramatic implications for any ideas concerning prenatal diagnosis and ‘prediction’ of any genotype/phenotype relationship (discussed more below).

Surprisingly, a precise definition of the term ‘oligogenic’ is not apparent or consistent in the world literature. Some people have invoked the term ‘oligogenic’ to mean an interaction between mutations in two genes to somehow collectively ‘cause’ a disease, such as with this above case of triallelism in Bardet-Biedl syndrome (Beales et al., 2003). These authors define oligogenic inheritance as occurring “when specific alleles at more than one locus affect a genetic trait by causing and/or modifying the severity and range of a phenotype” (Beales et al., 2003). Another case in point involves the 22q11.2 locus, also known as velocardiofacial syndrome. This deletion does not involve only a single gene, but rather ~X number of genes, depending on the exact size of the deletion interval. The phenotypic manifestations can be incredibly heterogeneous, illustrated by the fact that some ~30% develop psychotic symptoms and get labeled as ‘schizophrenic’ (Philip and Bassett, 2011). Of course, heuristic diagnoses for schizophrenia are usually made based on certain semantic criteria, so it is likely that subthreshold symptoms are not counted (or perhaps not even detected). But, at least one has the advantage of knowing which people possess the deletion, allowing one to perform detailed phenotyping to determine whether subthreshold symptoms were missed within a family, and this has indeed been done in the case of a well-known translocation involving *DISCI* (Blackwood et al., 2001; Hamshere et al., 2005). Unfortunately, genome-wide studies are not yet performed routinely for people with ‘idiopathic schizophrenia’, so it has been difficult to identify and group many people by genotype(s). As we discuss below, we believe that the routine clinical use of exome and eventually whole genome sequencing might finally enable this to occur, assuming that aggregation of genotype and phenotype data is allowed on a massive scale.

The definition of ‘polygenic’ literally means “many genes”, including the combined effects of dozens (or perhaps even hundreds) of different mutations in different genes on a particular phenotype, although it is sometimes not very clear whether these multiple genes are meant to be spread across individuals or within individuals. We tend to favor the definition involving multiple mutations within the same individual somehow contributing toward phenotypic development. Height has historically been characterized as being a polygenic phenotype, with GWAS studies implicating the possible involvement of hundreds of loci (Berndt et al., 2013; Visscher et al., 2010). Height is an easily measured phenotype and is generally described as being distributed continuously within human populations, modified of course by gender and ancestral backgrounds. If one looks at height in males or females of a certain ethnic background and from the same geographic locale, one can typically draw a semi-Gaussian (normal) function, but with tails that deviate from what is expected, encompassing rare cases of dwarfism and gigantism. We tend to also think that a single vertical measurement does not capture the true phenotypic variability involving height, as this measurement does not adequately capture the variability that exists in the many determinants of height (i.e., bone dimensions, age, environment, etc.). So, for a trait that seems conceptually simple to measure, there exists difficulty in uncovering its genetic component(s) due, in part, to uncharacterized uncertainty (variability) introduced at the phenotypic measurement level. If we now consider psychometrically defined traits, a large amount of further uncertainty is introduced at the phenotypic measurement level, as we are still unable to accurately characterize even a single measurement for most psychiatric disorders. These difficulties are underscored by the fact that psychiatric definitions are ephemeral and can change in a dramatic fashion over the course of even a few years. It seems premature to argue that schizophrenia, for example, is Gaussian in nature (Visscher et al., 2011). We would argue that we simply do not know enough about the phenotypic expression of the many different diseases that this amorphous concept of‘schizophrenia’ encompasses to be able to make any conclusions regarding its genetic inheritance on a population or individual level (Mitchell, 2012). Until there is substantial evidence to support another viewpoint, it is therefore important to treat each family as a special case. One must study people within families to determine whether some people in families have illness due to mutations with variable expression, modified by genetic background and environmental influences.

There have been numerous reviews concerning the ongoing debate for common and rare variants, with arguments made for various ‘camps’ of thought, including the common disease-common variant (CDCV) model, the infinitesimal model, the rare allele model and the broad sense heritability model (Gibson, 2011). Frankly, these models are simply semantic and reductionistic arguments that do not reflect the complexity of the human condition, and we are not sure that arguing for and against various models is useful, given that these models are basically straw men artificially constructed to be knocked down. This is very similar to the psychiatric literature in which several people decided, about 100 years ago, to introduce various names (or models) for certain diseases, such as the words ‘schizophrenia’ (Bleuler, 1958) and ‘manic-depressive illness or bipolar’ (Kraepelin, 1921). It is quite apparent to most clinicians that the phenotypic heterogeneity of these illnesses is so tremendous as to render these names basically moot and not particularly useful. This is akin to 50 years ago when people simply stated that someone had ‘cancer’. Now, it is not useful to say only that someone has cancer, as there are literally hundreds of molecular etiologies for cancer, divided up not only by organ expression but also by specific pathways in the cell (Sporn, 2011). We anticipate that in 50 years, these terms ‘schizophrenia’ and ‘bipolar’ will be replaced by much more precise molecularly defined terms, as is occurring now in the cancer field (Mukherjee, 2010; Vogelstein et al., 2013). Locus heterogeneity will likely play an important role in most diseases, but particularly in psychiatric disease, given the extensive phenotypic heterogeneity. Some of this complexity has been documented in reports of individual people (Eichenbaum, 2013; Luria, 1972,1976; Lyon, 2008; Lyon et al., 2008; Lyon and Coffey, 2009; Penrose, 1963; Ratiu et al., 2004; Sacks, 1995,1998; Van Horn et al., 2012; Ward, 1998; Worthey et al., 2011), and a review by one of us of the literature related to schizophrenia (Lyon et al., 2011) rendered the distinct impression that we really hardly know anything about the mechanistic basis of these many illnesses that we currently lump together as ‘schizophrenia’. This is primarily due to overly broad descriptions and categorizations of these illnesses into these artificially named syndromes, despite the obvious heterogeneous and inconsistent nature of these categorizations. Remarkably, bipolar and schizophrenia have been artificially ‘split’ into different syndromes (Craddock and Owen, 2010; Williams et al., 2011), in spite of the existence of a well documented literature demonstrating overlap in at least some families with symptoms from both ‘syndromes’ (Lichtenstein et al., 2009).

Oddly enough, some diseases such as Fragile X, Rett Syndrome and other now molecularly defined disorders are sometimes removed from the ‘nonsyndromic idiopathic autism’ camp, leaving the remaining disorders still eligible for a semantic debate about which ‘genetic model’ they fit into (Reiss, 2009). One wonders if the same thing has occurred for velocardiofacial syndrome, with its relevance to schizophrenia, given the overwhelming evidence that the single 22q11.2 deletion event predisposes its carriers to some version of ’schizophrenia’ with some exhibiting anywhere between 20 and 30% of the symptoms currently being defined as consistent with ‘schizophrenia’(Philip and Bassett, 2011). All of these disorders were at one point labeled as ‘idiopathic’ until molecular lesions associated with them were identified. It has been known by at least some researchers and clinicians for quite some time that there are likely many minor physical anomalies in people labeled as ‘nonsyndromic’ (Aldridge et al., 2011; Miles, 2011), all of which is further proof of the substantial phenotypic expression differences of all disorders. Therefore, the dichotomous use of the words ‘syndromic’ and ‘nonsyndromic’ is completely artificial and does not reflect the reality or complexity of the situation in any one person.

A recent paper using exome sequencing to study hypertension pedigrees made the following statements: “These findings demonstrate the utility of exome sequencing in disease gene identification despite the combined complexities of locus heterogeneity, mixed models of transmission and frequent de novo mutation. Gene identification was complicated by the combined effects of locus heterogeneity, two modes of transmission at one locus, and few informative meioses. Many so far unsolved Mendelian traits may have similar complexities. Use of control exomes as comparators for analysis of mutation burden may be broadly applicable to discovery of such loci “(Boyden et al., 2012). This paper illustrates exactly what we are discussing above, in terms of the possible heterogeneity of many illnesses on many levels, making it impossible to predict (or even need) any particular model that may or may not fit the disease. It is far better to allow the data to speak for themselves.

## De novo mutations, germline mosaicism and other complexities

Although this concept of somatic mosaicism has been in the literature for many years (Bakker et al., 1989; Hall, 1988; Hollander, 1975; Sastry et al., 1965; Vig, 1978), it is really only recently that more people are beginning to realize that it might be much more extensive in humans than previously thought (Baugher et al., 2013; Biesecker and Spinner, 2013; Choate et al., 2010; Coufal et al., 2011; Huisman et al., 2013; Jongmans et al., 2012; Kurek et al., 2012; Lindhurst et al., 2012; Lindhurst et al., 2011; Lyon and Wang, 2012; Macosko and McCarroll, 2012; Margari et al., 2013; Shirley et al., 2013; Steinbusch et al., 2013; Tanaka et al., 2012; Weiss, 2005; Yamada et al., 2012). In fact, hardly anything is truly known regarding the extent of somatic mosaicism in humans and its effect on phenotype in even well studied diseases. For example, little is known regarding pathogenesis of the phenotype in people with trisomy 21 mosaicism and Down syndrome, although there is likely variation in phenotype associated with the percentage of trisomic cells and their tissue-specificity (Hulten et al., 2010; Iourov et al., 2008; Kovaleva, 2010). A more recent study looked at this issue of somatic mosaicism in Timothy syndrome type 1 (TS-1), which is a rare disorder that affects multiple organ systems and has a high incidence of sudden death due to profound QT prolongation and resultant ventricular arrhythmias. All previously described cases of TS-1 are associated with a missense mutation in exon 8A (p.G406R) of the L-type calcium channel gene (Ca(v)1.2, *CACNA1C*). Most cases reported in the literature represent highly affected people who present early in life with severe cardiac and neurological manifestations, but these authors found somatic mosaicism in people with TS-1 with less severe manifestations than the typical person with TS-1 (Etheridge et al., 2011). There are therefore likely large ascertainment biases, given that people with subtler phenotypes are likely not coming to anyone’s attention. The implications of these findings with somatic mosaicism are that one cannot currently predict phenotype from genotype, particularly in the absence of any comprehensive characterization of which tissues are mutated in any one person. Also, putative ’de novo’ mutations can instead represent cases of parental mosaicism (including in the germline), which could be revealed by careful genotyping of parental tissues other than peripheral blood lymphocytes. In fact, we are increasingly becoming aware of many instances of germline mosaicism, in which a mutation is not present or is present only at a very low level in the blood sample from a parent, but clearly must be in their germline, as they have two or more children with the same mutation that must therefore have originated through the parent’s germline (Aldred et al., 2000; Barbosa et al., 2008; Chaturvedi et al., 2000; Evans et al., 2006; Frank and Happle, 2007; Hosoki et al., 2005; Jongmans et al., 2008; Mari et al., 2005; Meyer et al., 2012b; Parodi et al., 2008; Pauli etal., 2009; Rand et al., 2012; Sato et al., 2006; Sbidian et al., 2010; Shanske et al., 2012; Slavin et al., 2012; Sol-Church et al., 2009; Tajir et al., 2013; Trevisson et al., 2014; Venancio et al., 2007; Wuyts et al., 2005). Clearly, we are truly ignorant concerning the extent of diversity brought about by somatic mosaicism, and it is therefore far too simplistic to assume that a single blood draw truly represents the entire genome of a human being, with anywhere from 25-100 trillion cells in their body divided up among multiple organs and other tissue systems. Of course, even the words “whole genome sequencing” are misleading, as there might very well be millions to trillions of similar (but not the exact same) genomes in each person’s body.

## Rare and compensatory mutations

There is an increasingly rich literature regarding rare mutations with seemingly large phenotypic effects (Boyden et al., 2002; Jonsson et al., 2012; Styrkarsdottir et al., 2013; Williams, 2004). An example of this is Liam Hoekstra, known as the world’s strongest toddler when he was age 3, and who has an extremely rare mutation in the gene encoding myostation, leading to myostatin-related muscle hypertrophy with increased muscle mass and reduced body fat. However, the effects of these mutations have mainly been reported in the context of particular genetic backgrounds, and so our knowledge of the expression of these mutations in the context of any number of genetic backgrounds is lacking. It is likely that there can be, and are, many genomic elements that act in concert to influence these traits in a phenotypic spectrum. Of course, compensatory mutations can be explored in the context of other organisms (Esvelt et al., 2011; Fu et al., 2013a; Leconte et al., 2013), but human migration and breeding is certainly not something that can be experimentally manipulated!

There are many disabling psychiatric syndromes, which have been lumped under certain artificial categories, such as schizophrenia, Tourette Syndrome (TS), obsessive compulsive disorder (OCD), and attention deficit hyperactivity disorder (ADHD). A very good way forward is to study these syndromes in large families living in the same geographic region, so as to control for ancestry differences, minimize environmental influences, and focus on specific genotypes in these families. It is possible that a low number of genetic mutations will be shared in a relatively small combination (on the order of 1-3 such variants) among affected relatives within some pedigrees, and that these variants will not be present in the same combination in unaffected relatives or in other families with very little to no neuropsychiatric disorders (Crepel et al., 2010; Fullston et al., 2011; Girirajan et al., 2012; Lyon and Wang, 2012; Mitchell, 2012; Mitchell and Porteous, 2011; Shi et al., 2013). An alternative is that some affected people in these families have these illnesses due to additive and/or epistatic interactions among dozens to hundreds of loci within each person (Klei et al., 2012; Zuk et al., 2012). The currently classified syndromes of schizophrenia, obsessive compulsive disorder (OCD), attention deficit hyperactivity disorder (ADHD), autism and other mental illnesses are quite heterogeneous within and between families, and these symptoms have also been observed in known single locus disorders such as Fragile X and 22q11.2 velocardiofacial syndrome (Girirajan et al., 2012; Mitchell, 2012).

Some of these syndromes are referred to as ‘complex’ diseases simply because the presentation is so incredibly heterogeneous that is it very likely that there will be multiple different genetic and environmental explanations. One possible genetic explanation is that some symptoms of severe mental illness may emerge in a particular family due to a genetic constellation including dozens to hundreds of loci acting in each person either additively or via epistasis (and possibly modified by environment; G X E), which some refer to as the ‘polygenic’ model (Anney et al., 2012; Klei et al., 2012; Visscher et al., 2011; Zuk et al., 2012), as previously discussed. If true, for predictive efforts in any particular family, the solution will ultimately require whole genome sequencing to tease out the numerous mutations involved. On the other hand, some discuss this concept of “many rare variants of large effect”, which they refer to as the ‘oligogenic’ model of inheritance (Gagnon et al., 2011; Schaaf et al., 2011), as previously discussed. Some families have deleterious copy number variants (Elia et al., 2010; Gai et al., 2012; Girirajan et al., 2012; Malhotra and Sebat, 2012; Shaikh et al., 2011), and de novo single nucleotide mutations have recently been implicated as important for spontaneous ‘singleton’ cases in at least some families (Iossifov et al., 2012; Neale et al., 2012; Novarino et al., 2012; O’Roak et al., 2012b; Sanders et al., 2012; Xu et al., 2012). There could also be a set of families with single, pair or triplet interactions among 1-3 gene mutations of high expression that can largely, on their own, contribute to a set of symptoms currently overlapping with named syndromes, such as ‘autism’ and ‘schizophrenia’ (Girirajan et al., 2010). As there is no way of really distinguishing between these two artificially created models in any one particular family, it is reasonable (with current costs) to perform microarray genotyping and whole genome sequencing as a comprehensive way to ascertain most of the relevant genetic variance in any particular family.

It is becoming generally accepted that at least 5% of the ‘autisms’ appear to be associated with various large copy number variants (Sanders et al., 2011). So, it is likely that some additional portion of the ‘autisms’ will be influenced by other types of mutations, with some evidence pointing to a role for ‘de novo’ mutations in singleton, uninherited cases of autism (Iossifov et al., 2012; Neale et al., 2012; O’Roak et al., 2012a; O’Roak et al., 2012b; Sanders et al., 2012) and other evidence suggesting that there might be multiple genetic and environmental influences in each person (Klei et al., 2012).

## Current ability / approaches

There has been an explosive growth in exome and whole genome sequencing (WGS) (Lyon and Wang, 2012), led in part by dramatic cost reductions. The same is true for genotyping microarrays, which are becoming increasingly denser with various markers while maintaining a relatively stable cost (lllumina, 2013). With rapid advancements in sequencing technologies (Schneider and Dekker, 2012) and improved haplotype-phasing (Peters et al., 2012; Williams et al., 2012), high-throughput sequencing (HTS) data on the genomes of a diverse number of species are being generated at an unprecedented rate. The development of bioinformatics tools for handling these data has been somewhat lagged in response, creating a gap between the massive data being generated, and the ability to fully exploit their biological content. Many short read alignment software tools are now available, along with several single nucleotide variants (SNVs) and copy number variant (CNVs) calling algorithms (Lyon and Wang, 2012). However, there is a paucity of methods that can simultaneously handle a large number of genetic variants and annotate their functional impacts (particularly for a human genome, which typically hosts >3 million variants), despite the fact that this is an important task in many sequencing applications. Functional interpretation of genetic variants therefore becomes one of the major obstacles to connect sequencing data with biomedical researchers who are willing to embrace the sequencing technology.

In the medical world, WGS has since led to the discovery of the genetic basis of Miller Syndrome (Roach et al., 2010) and in another instance, it was used to investigate the genetic basis of Charcot-Marie-Tooth neuropathy (Lupski et al., 2010), alongside a discussion of the ‘return of results’ (McGuire and Lupski, 2010). In 2011, the diagnosis of a pair of twins with dopa (3,4-dihydroxyphenylalanine) responsive dystonia (DRD; OMIM #128230) and the discovery that they carried compound heterozygous mutations in the SPR gene encoding sepiapterin reductase led to supplementation of 1-dopa therapy with 5-hydroxytryptophan, a serotonin precursor, resulting in clinical improvements in both twins (Bainbridge et al., 2011).

Despite current technological limitations, mutations are continually being identified in research settings (Bamshad et al., 2011; Hedges et al., 2009; Lyon, 2011; Ng et al., 2010a; Ng et al., 2010b; Roach et al., 2010). However, the human genomics community has recognized a number of distinct challenges, including with phenotyping, sample collection, sequencing strategies, bioinformatics analysis, biological validation of variant function, clinical interpretation and validity of variant data, and delivery of genomic information to various constituents (Katsanis and Katsanis, 2013; Lyon and Wang, 2012). In particular, there is a need for large pedigree sample collection, high-quality sequencing data acquisition, rigorous generation of variant calls, and comprehensive functional annotation of variants (Lyon and Wang, 2012). Empirical estimates seem to suggest that exome sequencing can identify a putative disease associated variant in only about 10-50% of the cases for which it is applied (Lyon and Wang, 2012), and the genetic architecture of most neuropsychiatric illness is still largely undefined and controversial (Klei et al., 2012; Mitchell, 2012; Mitchell and Porteous, 2011; Visscher et al., 2011). The sequencing of entire genomes in large families will create a dataset that can be analysed and re-analysed for years to come as new biology and new methods emerge. The cost of a whole genome will likely decrease much more rapidly in relation to the cost of exome sequencing, given the relatively fixed labor and reagent costs for capturing the exons in the genome. Also, there is emerging evidence that exon capture and sequencing only achieves high depth of sequencing coverage in about 90% of the exons, whereas WGS does not involve a capture step and thus obtains better coverage on >95% of all exons in the genome. Of course, even the definition of the exome is a moving target, as the research community is constantly annotating and finding new exons not previously discovered (Wu et al., 2013; Zumbo and Mason, 2014), and therefore WGS is a much more comprehensive way to assess coding and non-coding regions of the genome.

It is obvious that in both research and clinical settings WGS can dramatically impact clinical care, and it is now a matter of economics and feasibility in terms of WGS being adopted widely in a clinical setting (Lyon, 2012; Lyon and Wang, 2012). There are, however, still many challenges in showing how any one mutation can contribute toward a clear phenotype, particularly in the context of genetic background and possible environmental influences (Moreno-De-Luca et al., 2013b). Bioinformatics confounders, such as poor data quality (Nielsen et al., 2011), sequence inaccuracy, and variation introduced by different methodological approaches (O’Rawe et al., 2013) can further complicate biological and genetic inferences. Furthermore, one cannot exclude polygenic and epistatic modes of inheritance (Bloom et al., 2013; Davis et al., 2011; El-Hattab et al., 2010; Kajiwara etal., 1994; Katsanis et al., 2001; Lai et al., 2010; Lemmers et al., 2012). To address these issues, future work will need to focus on evaluating next generation sequencing data coming from multiple sequencing and informatics platforms, and involving multiple other family members. By using a combination of data from many family members and from different sequencing technologies evaluated by a number of bioinformatics pipelines, we can maximize accuracy and thus the biological inference stemming from these data.

## Prenatal diagnosis, preimplantation genetic diagnosis/screening

> *“Before a new function can arise, it may be essential for a lineage to evolve a potentiating genetic background that allows the actualizing mutation to occur or the new function to be expressed. Finally, novel functions often emerge in rudimentary forms that must be refined to exploit the ecological opportunities. This three-step process—in which potentiation makes a trait possible, actualization makes the trait manifest, and refinement makes it effective—is likely typical of many new functions*.” —*<u>Richard Lenski</u> (Blount et al, 2012)*

A great clinical geneticist, John Opitz, has observed the following: “More fetuses die prenatally than are born alive. Many die because of genetic conditions, malformations, and syndromes. Most are not autopsied, and in such cases appropriate genetic counseling is not provided or possible. In such ‘cases’ (fetuses, infants) a huge amount of genetic pathology is yet to be discovered (our last frontier!)” (Opitz, 2012).

In this regard, some have suggested a canalization model, which describes phenotypes as being robust to small perturbations, seemingly stuck within “phenotypic canals”. Phenotypes may ‘slosh’ against the sides of the canal during development, but with little effect on the final outcome of development (Waddington, 1959, 2012). In such a model, it is only perturbations with a magnitude exceeding a certain threshold that can direct the developmental path out of the canal (see Figure 1 for an illustrative model of canalization). Accordingly, phenotypes are robust up to a limit, with little robustness beyond this limit. This pattern may increase rates of evolution in fluctuating environments, as phenotypes are more likely to be perturbed with increased frequency and magnitude, thus leading to more rapid delineations and differentiations of canalized phenotypes.

One could argue that the birth of a child in one particular famliy with a clear phenotype, such as cystic fibrosis, along with previously identified associated mutations, dramatically increases the ‘prior probability’ that a future child with these same mutations being born in that same family would have a similar ‘canalized’ phenotype. It is really only in that particular situation in which one could make a somewhat informed prediction of genotype going down one particular phenotypic “canal”. And yet, a study in Australia from 2000-2004 showed that of the 82 children born with cystic fibrosis (CF) in Victoria, Australia, 5 (6%) were from families with a known history of CF. The authors found that “even when a family history is known, most relatives do not undertake carrier testing. In an audit of cascade carrier testing after a diagnosis of CF through newborn screening, only 11.8% of eligible (non-parent) (82/716) relatives were tested” (McClaren et al., 2011). These same researchers also showed that in a clinical setting, the diagnosis of a baby with CF by newborn screening “does not lead to carrier testing for the majority of the baby’s non-parent relatives” (McClaren et al., 2010). This is incredibly unfortunate, given that predictions of any reliability ought to include the prior probability of someone being born in that ‘ancestry group’ with the mutations and phenotype of interest.

Despite the above facts, non-invasive sequencing of fetal genomes is an area of intense interest in genomic medicine, and a cynical person might argue that the rush to implement this technology is driven mainly by financial interests. Current techniques are based on the observation that a small proportion of the cell-free DNA in a pregnant woman’s blood is derived from the fetus, so that aneuploidy or genomic sequence of a fetus may be inferred by sequencing of maternal plasma DNA and algorithmic decoupling of maternal and fetal DNA variants. A few companies are already marketing non-invasive prenatal screening (NIPS) tests for non-invasive detection of trisomy 21 associated with Down’s syndrome. One can reasonably argue that detecting Down’s syndrome is a conceptually and practically much simpler task than detecting individual variants within the fetal genome to assess mutations associated with disorders such as cystic fibrosis and hearing loss. However, with sufficiently high sequence depth, it is technically feasible to detect single nucleotide alterations in a fetal genome, as shown in several recent papers (Cheng et al., 2013; Fan et al., 2012; Kitzman et al., 2012; Papageorgiou and Patsalis, 2013). But, to allow accurate detection of individual variants, very high sequencing depth is required (potentially hundreds-fold higher than sequencing germline genomes); therefore, it is likely that targeted exon capture and sequencing might dominate the market until sufficiently high depth whole-genome sequencing becomes an economically feasible alternative. Given these technological developments, it is likely that some form of fetal genome testing will be available in the next few years. Others have noted that we might be reaching a point in the near-term future where it may be feasible to incorporate genetic, genomic and transcriptomic data to develop new approaches to fetal treatment (Bianchi, 2012; Guedj and Bianchi, 2013). One concern is that greed and financial conflicts of interest could lead to indiscriminate marketing and use of NIPS as diagnostic tests, rather than simply as screening, and that this technology will be implemented without any regard for genetic background or environmental differences, alongside a complete misunderstanding of this concept of extreme variability in phenotypic expression.

## Implications for acceptance, prognosis and treatment

> *“When a complex system starts to dysfunction, it is generally best to fix it early. The alternative often means delaying until the system has degenerated into a disorganized, chaotic mess* — *at which point it may be beyond repair. Unfortunately, the general approach to cancer has ignored such common sense. The vast majority of cancer research is devoted to finding cures, rather than finding new ways to prevent disease”* —*<u>Michael Sporn</u> (Sporn, 2011).*

Prevention of illness through environmental modification has been, and likely always will be, the major driver for global health (Mukherjee, 2010; Sporn, 2011). With this in mind, the sequencing of whole genomes on a large scale promises to enable the discovery and prediction of disease in some people. The ability to sequence an infant at birth and to be able to predict a higher probability of certain phenotypes, such as developmental delay, would allow for educational and behavioral interventions to influence the phenotype, thus
altering the trajectory of that phenotype (Bates et al., 2014; McIntyre, 2008; Rickards et al., 2007, 2009; Velleman and Mervis, 2011). One recent study of chromosomal microarray (CMA) testing found that “among 1792 patients with developmental delay (DD), intellectual disability (ID), multiple congenital anomalies (MCA), and/or autism spectrum disorders (ASD), 13.1% had clinically relevant results, either abnormal (n = 131; 7.3%) or variants of possible significance (VPS; n = 104; 5.8%). Abnormal variants generated a higher rate of recommendation for clinical action (54%) compared with VPS (34%; Fisher exact test, P = 0.01)” (Coulter et al., 2011). The authors concluded that “CMA results influenced medical management in a majority of patients with abnormal variants and a substantial proportion of those with VPS” thus supporting the use of CMA in this population (Coulter et al., 2011). We agree that the identification of certain CNVs and other mutations can suggest a range of phenotypes that might occur in any one individual with that mutation or mutations.

However, there are some major barriers to the widespread implementation of genomic medicine in the clinic. These include:

1. Lack of public education
2. Lack of physician knowledge about genetics
3. Apathy on the part of the populace in terms of preventive efforts
4. Refusal of insurance companies and governments to pay for genetic testing
5. Focus in our society on treatment, not on early diagnosis and prevention
6. Privacy concerns
7) Limits of our current knowledge

The emphasis should be on diagnosis and prevention, not just on treatment. During the medical training of one of the authors (GJL), two episodes helped to illustrate this. The first involved a 15-year old girl with Type I diabetes, who was hospitalized dozens of times with diabetic ketoacidosis. Literally hundreds of thousands of dollars were spent to repeatedly save her life, but very little time or money was spent on therapy or education to teach her about taking her insulin and ensuring that she did. Unfortunately, in America at least, this is due to a relative lack of reimbursement for such activities, whereas saving someone already in diabetic ketoacidosis is quite lucrative to everyone involved. A second episode involved a 14-year old boy, who had been hospitalized well over 10 times with acute pancreatitis over a ten year period, with very little thought concerning why he had recurring pancreatitis. Finally, someone obtained a genetics consult, and they recommended cystic fibrosis (CF) genetic screening, which had never been ordered before due to a prior ‘negative’ sweat test. It turns out that this boy had two rare mutations in *CFTR,* undiagnosed till then, which had been contributing to recurrent pancreatitis. He had never had any lung manifestations, and he had never had a positive sweat test for CF, mainly due to the fact that these mutations appeared to only be exerting effects in his pancreas, not in his skin or lungs. After this diagnosis, this person benefited from pancreatic enzyme supplementation, along with therapy and education. Once again, the reason it took so long to diagnose this person is because the incentive structure in many developed nations is not on early diagnosis and prevention, but rather on treatment of people only once they become severely ill (Brawley and Goldberg, 2012; Makary, 2012). This is illustrated by the fact that there are only about ~1000 medical geneticists in America and ~3000 genetic counselors, for a population of~315 million, which makes it basically impossible for these limited number of professionals to implement genomic medicine in any meaningful way (Brandt et al., 2013). The numbers of such health care professionals are even smaller in developing regions of the world, thus making it currently very difficult to provide widespread genetic counseling (Bittles, 2013; Bittles and Black, 2010; Hamamy, 2012). Stepping into this void are direct-to-consumer for-profit genetic testing companies, and this is certainly one disruptive way of trying to help people manage their genetic results online (Chua and Kennedy, 2012; Francke et al., 2013), although financial motives and lack of transparency can create problems (Sterckx et al., 2013).

Privacy concerns have added to the difficulties of implementing genomics-guided medicine. Genetic data have the potential of being informative across a wide variety of human traits and health conditions, and some worry about the potential misuse of these data by insurance agencies as well as by health care providers (Allain et al., 2012). Genetic testing has historically been focused on targeting and examining a small number of known genetic aberrations (Bakker, 2006); however, since the advent of high-throughput sequencing technologies, the landscape is starting to change. With the emergence of tests that can target and examine all coding regions of the genome, or even the genome in its entirety, testing can now be performed on a more global and exploratory scale. Some people worry about returning the results of such a test, whose findings can have questionable clinical significance, and in response have advocated for selectively restricting the returnable medical content. Others have proposed complicated anonymization techniques that could allow for a safe return of research results to participants whose genome is suspect to contain ‘clinically actionable’ information. One such proposition involves the cryptographic transformation of genomic data in which only by the coalescence of keys held by many different intermediate parties would the identity of the participant be revealed, and only in cases where all parties agree that there is indeed the presence of clinically actionable information (Hunter et al., 2012). These types of recommendations take a more paternalistic approach in returning test results to people, and generally involve a deciding body of people that can range in size from a single medical practitioner to a committee of experts. In contrast, there is a growing movement among the populace to learn more about their own ‘personalized’ health and health care. There has also been a renewed push for the unfiltered sharing and networking of health related data, which has been facilitated and hastened by the explosion of digitally mediated social networking over the past decade, as well as by efforts from 23andMe (Kranhold, 2007) and the Personal Genomes Project (Ball et al., 2012) that aim to popularize and democratize genetic testing. Clearly, between these contrasting approaches, there is a tradeoff between the privacy and personal safety one can expect to retain by either freely acquiring and sharing the full breadth of one’s genetic testing data, or by allowing deciding bodies to choose what information you will receive.

Public databases containing human sequence data have grown in magnitude and in number, and relatively comprehensive sequencing data have already been generated and published on thousands of people (Abecasis et al., 2012; Fu et al., 2013b). Similar privacy concerns have since been expressed about the degree of medical and personal privacy that these and other research participants can expect (McGuire and Gibbs, 2006), given that each person is genetically unique. As a demonstration of current vulnerabilities, researchers have shown that the identities of participants can be discovered using these publicly available data (Gymrek et al., 2013). Although these data have been instrumental in furthering our understanding of human genetics, medicine, and biological processes in general, some advocate for caution when sharing and publishing human genetic sequence information (Lowrance and Collins, 2007).

As the cost and difficulty of sequencing continually decreases, a wealth of data are becoming available to researchers, privately funded institutions and individual consumers. More people are willing to share a larger portion of their personal life in the public arena, and we fully expect that, given the popularization of ‘personalized’ genomic health related data, more people will want to share these data and offer their own DNA sequence for others to explore. There is a trade off between the risks inherent in sharing vast quantities of health data, and maintaining personal privacy in the burgeoning age of personalized medicine and genomics. As the technology and science mature, our power to interpret and use these health data for practical and preventative measures will certainly improve. Conventions for privacy and autonomy will likely be driven by popular demand, and could vary from person to person, as all people differ in their desire for privacy and autonomy (see Figure 2 for a conceptual model of this tradeoff).

In addition, within the current paradigm of genetic determinism, which stretches back to the time of William Bateson (Radick, 2005, 2013), some people would have us believe that variants can and should be binned into different classes based on clinical utility and validity (Berg et al., 2013; Goddard et al., 2013; Green et al., 2012), without any obvious regard to genetic background or environmental differences. Environment and ancestry matter (Radick, 2005, 2011, 2013; Weldon, 1902), and yet some clinical geneticists trained in the current paradigm of genetic determinism clearly do not wish to acknowledge this. Categorical thinking misses complexity. In fact, one medical academy in America recently released guidelines in which they recommended the “return of secondary findings” for only 57 genes, without any real guidance for the rest of the genome or environmental influences (Green et al., 2013). This is therefore a very conservative set of recommendations, given that there are approximately 20,000 protein-coding genes in the human genome, along with the thousands of other identified, important noncoding elements of the genome (Batista and Chang, 2013; Cartault et al., 2012; Hansen et al., 2013; Kapusta et al., 2013; Khoddami and Cairns, 2013; Ledford, 2013; Maxmen, 2013; Memczak et al., 2013; Mercer and Mattick, 2013; Miura et al., 2013; Moreau et al., 2013; Ning et al., 2013; Perrat et al., 2013; Sabin et al., 2013; Salzman et al., 2012; Wilusz and Sharp, 2013)! As stated above, but worth repeating, there are ~12 billion nucleotides of DNA in every cell of the human body, and there are 25-100 trillion cells in each human body. Given genetic modifiers, somatic mosaicism, epigenetic changes, and environmental differences, no two human beings are the same, and therefore the expression of any mutation will be different in each person. At best, phenotypes will follow canalized pathways in direct relatives, such as mother and child, so the analysis of mutations over several generations in the same families is a worthwhile effort. But, how we will ever get to a world of millions of whole genomes shared and analyzed for numerous additive, epistatic interactions and gene by environment interactions, so that we can make any reliable predictions for any one human being, if we are only recommending ‘return of results’ from ~57 genes? We need to sequence and collate online the raw exome and genome data and phenotypic information from thousands and then millions of people, so that we can actually begin to really understand the expression patterns of any mutation in the human genome in particular families. In medicine, people tend to create illusions of certainty, when in fact everything is probabilistic (Gigerenzer, 2002). Some humans like to be told things in a ‘yes/no’ manner, but there always exists a degree of unresolvable uncertainty.

## Conclusions

> *“A new scientific truth does not triumph by convincing its opponents and making them see the light, but rather because its opponents eventually die, and a new generation grows up that is familiar with it.”* — *<u>Max Planck</u>*

With the advent of exome and whole genome sequencing, we need to focus again on families over several generations, so as to attempt to minimize genetic differences, locus heterogeneity and environmental influences. Forging strong ties with families will also enable access to other tissues to continue to study newly discovered loci with many emerging technologies. Some might consider it to be ‘social activism’ to advocate for a more comprehensive collection and collation of human pedigrees, whole genome sequencing data and phenotypic information. But, in the words of one author: “Scientists, whether we like it or not, are members of society, and we are prone to the ideas and beliefs of the times in which we live (Mole, 2006).” We currently live within a paradigm of genetic determinism, but we should not be forever condemned to this simplistic mode of thinking. One can imagine or hope that in the not too distant future, each person will be able to keep track of detailed longitudinal phenotyping data on themselves online, and they will be able to link this to records of their relatives, both living and deceased. One can also hope that we are approaching a time where sufficient information is available within many large families for calculating highly accurate probabilistic outcomes (Gigerenzer, 2002; Gigerenzer and Galesic, 2012; Gigerenzer et al., 2010; Kurz-Milcke et al., 2008; Sokal, 2012), at which point we might be able to more effectively alter the trajectory for many diseases. One can see this beginning already to occur in certain geographically isolated clans, such as in Iceland (Jonsson et al., 2012; Styrkarsdottir et al., 2013), so there is some optimism that this can indeed occur on a global level, including in the currently less developed regions of the world (Bittles, 2013).

**Figure 1.**
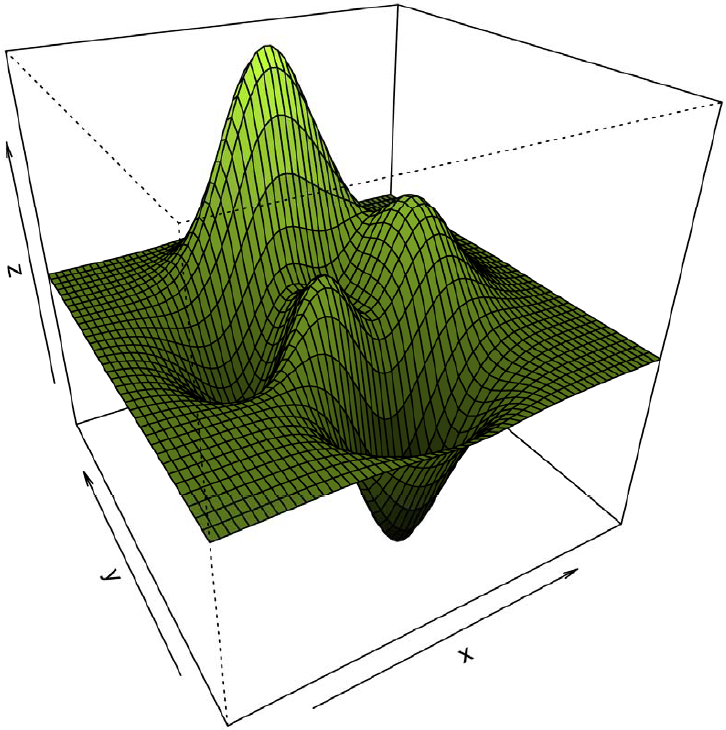
A conceptual model of canalization. The *y* plane represents a phenotypic spectrum, the *x* plane represents the canalized progression of development through time, and the *z* plane represents environmental fluctuations. As any particular phenotype progresses through development, it can encounter environmental fluctuations that either repel (a local maximum) or attract (a local minimum) its developmental path. Either force, if strong enough, can cause a shift in the developmental path, fundamentally altering the end resulting phenotype.

**Figure 2.**
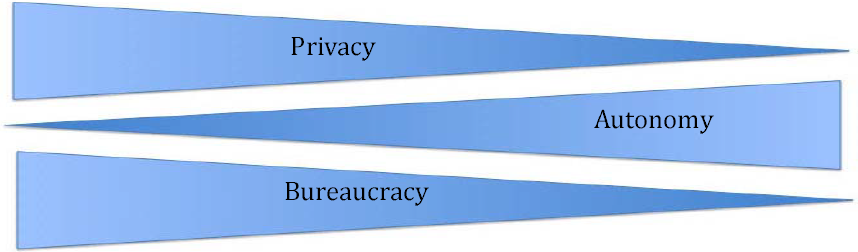
An illustration of the tradeoff between privacy and autonomy when receiving results from genetic testing. Models that guarantee an increased level of privacy are generally accompanied by a great deal of bureaucratic and paternalistic decision-making on the part of medical and advisory institutions (left). Models that propose and advocate for increased autonomy when receiving genetic test results come with the risk of reduced privacy (right). A whole genome sequence from a single person could, in principle, inform many aspects of his/her health care as well as allow for the prospect of future health predictions. This leads to speculations on how insurance agencies and health care providers could/would use this information. One can envision a ‘sinister scenario’ where people are rejected from hospitals and denied insurance based on putative genetic aberrations that may associate with costly, long term, care. Others worry about the potential implications of results found by genome scale testing, and would rather not know about risks pertaining to untreatable illnesses. Recent movements push for the democratization as well as large-scale adoption of this type of testing for every person, which could help to prove that we are all truly genetically unique and all carry any number of mutations and/or large genetic aberrations that may or may not be associated with disease. In reality, current technologies are far from the realm of genotype to phenotype predictions, and so genetic discrimination could only create illusory economic gains for any institution for the foreseeable future.

## Acknowledgements

We thank the editor, Kevin Mitchell, for detailed comments and suggestions regarding an earlier draft of this manuscript. Others who have made very helpful suggestions include: Anne Buchanan, George Church, Nathaniel Comfort, Jesse Gillis, Nathaniel Pearson, William Provine, Gregory Radick, Michael Stone, and Kai Wang. We also acknowledge members of the Lyon laboratory, particularly Han Fang and Max Doerfel, for their helpful suggestions as well. We would also like to thank Gail Sherman at the CSHL library for her efforts to procure some of the older literature on our behalf.

